# Slow diffusion and signal amplification on membranes regulated by phospholipase D

**DOI:** 10.1101/2024.07.08.602473

**Authors:** Gen Honda, Chihuku Tanaka, Satoshi Sawai, Miho Yanagisawa

## Abstract

Control of molecular diffusion is pivotal for highly fluidic membranes to serve as substrates for biochemical reactions and the self-assembly of molecular machinery driving membrane protrusions. Lateral diffusion in membranes depends on lipid composition, which is highly diverse and homeostatically controlled in living cells. Due to the complexity of the underlying processes, its impact on molecular diffusion remain largely unclear. In this study, we show that lipid diffusion in model membranes is markedly decreased in cytosolic extracts. The reduction in lipid diffusivity could be pharmacologically inhibited by targeting phospholipase D (PLD), and addition of PLD to membranes mimicked the reduction in diffusion. Phosphatidic acid, a direct product of PLD, diffused slowly in model membranes and reduced the diffusivity of surrounding lipids. Furthermore, we demonstrated that PLD specifically controls the lateral diffusion of a myristoylated protein in cells, possibly due to auxiliary electrostatic interactions between cationic residues located near the lipidated tail and anionic phospholipids. PLD controlled the size and lifetime of localized patches of phosphatidylinositol (3,4,5) triphosphates that specify regions of membrane protrusions. Overall, the results of this study suggest that PLD controls the lateral diffusion of certain membrane proteins, which play key roles in phosphoinositide signaling.

**Significance Statement:** In living cells, many biochemical reactions occur in confined regions on the membranes, facilitating the local occurrence of specific events, such as membrane protrusion. This is puzzling from a physical perspective because the membrane is a two-dimensional fluidic structure that should allow molecules to spread freely. Herein, we found that the fluidity of artificial membranes was markedly reduced by adding extracts from the cell cytoplasm. A lipid-modifying protein phospholipase D (PLD) was found to be responsible for this and it regulates the diffusion of membrane proteins in cells. This study suggests the novel role of PLD as a regulator of molecular diffusion and its impact on phosphoinositide production that serves as an important signal for cell deformation.

## Introduction

The cell membrane serves as a scaffold for plenty of biochemical processes. Transmembrane receptors, small GTPases, and phospholipids move around within the membranes, triggering signaling cascades upon stochastic collisions. Given the known diffusion constants of lipids and proteins in the classical fluid mosaic model (1), it is not trivial how enzymatic reactions can outpace the attenuation due to molecular diffusion (2). According to single-particle tracking (SPT) measurements, membrane proteins undergo multimodal diffusion (3–6). Various hidden structures have been proposed to explain the constraints on lateral diffusion. For instance, dynamically segregated domains of certain lipids and proteins may act as membrane compartments. The liquid-ordered phase, referred to as a lipid raft, consists of saturated fatty acids tightly packed with sphingolipids and sterols. Membrane proteins acylated with myristate or palmitate and glycosylphosphatidylinositol (GPI)-anchored proteins are partitioned into rafts, which are assumed to serve as signaling platforms (7). In addition, F-actin meshwork adjacent to the membrane has been reported to impede molecular diffusion (6, 8–10). For example, cortical actin limits the diffusion of Fc-gamma receptors at the trailing edge of macrophages (6). Adhesion receptors such as integrins and cadherins are assembled into clusters stabilized by F-actin with the help of force-sensitive molecules such as talin and vinculin (5, 11). However, it should be noted that local signaling events can occur on the plasma membrane independent of the actin network. A prominent example is the formation of micron-sized patches rich in activated Ras GTPase and phosphatidylinositol (3,4,5)-trisphosphate (PI(3,4,5)P3), which specify membrane domains that subsequently become the macropinocytic and phagocytic cups (3, 12–17). The signaling patches are known to form and travel in actin-free membranes (14) and are characterized by the depletion of membrane-anchored proteins due to high diffusion rates in the patch region (3). Despite their importance in cell polarity, the origin of the difference in the diffusion rates that induces macroscopic segregation remains unclear.

Lateral diffusion in lipid membranes is primarily determined by lipid packing density and specific molecule–lipid interactions. Diverse lipid species with different combinations of head groups and fatty acid unsaturation are produced in cells. Lipid composition is homeostatically controlled by metabolism and vesicle trafficking to maintain a nearly constant membrane fluidity (18, 19). Therefore, reconstitution approaches are crucial for visualizing changes in membrane fluidity by eliminating associated homeostatic effects. Previous studies using cell extracts have reported the irreversible progression of biochemical reactions outside living cells, which emerged in unusual forms such as extraordinarily long actin filaments (20) and short-lived Rho-actin waves reconstituted *in vitro* (21). Unlike in live cells, the cytosolic extracts used in these studies facilitated lipid partitioning (20, 22) and reduced lipid diffusivity (21) in solid-supported membranes. However, the molecular entities underlying these phenomena remain unknown. In this study, we investigated the fluidity of supported lipid bilayers in the presence of *Dictyostelium* cell extracts. We implicate phospholipase D (PLD) as a key factor in reducing lipid diffusivity and demonstrate that PLD controls the lateral diffusion of myristoylated kinase in living cells. The results of this study suggest that PLD-induced slow diffusion is pivotal for regulating the size, lifetime, and magnitude of phosphoinositide signaling.

## Results

### Cell extracts reduced the fluidity of solid-supported model membranes

To explore the mechanisms controlling diffusion in cell membranes, we investigated the effect of cell extracts to alter the fluidity of model lipid membranes. Phosphatidylcholine (PC) is one of the most abundant phospholipids in eukaryotic cells, accounting for approximately 30% (23). Therefore, we performed fluorescence recovery after photobleaching (FRAP) analysis using supported lipid bilayers (SLBs) consisting of 1,2-dioleoyl-sn-glycero-3-PC (DOPC) and 1,2-dimyristoyl-sn-glycero-3-phosphatidylethanolamine bound to lissamine rhodamine B (Rh-DMPE). SLBs were formed on borosilicate coverslips using the vesicle fusion method, and fresh cytosolic extracts of *Dictyostelium discoideum* cells were prepared using the French press cell lysis method. Within the first 30 min after applying the cell extract, numerous dark regions appeared in the SLB, which were likely due to lack of lipids or phase separation (Movie S1). Similar observations have been reported for *Dictyostelium* (24) and *Xenopus* egg extracts (20). In phosphate buffer (PB) controls, the fluorescence of Rh-DMPE remained uniform for at least 5 h. After bleaching, the fluorescence recovered within seconds (Fig. 1A, “PB” and Fig. 1B, “PB”); the estimated diffusion coefficient of 0.74 ± 0.08 µm^2^/s is within the typical range known for lipids (7). In contrast, the fluorescence showed no recovery in cell extracts (Fig. 1A, “Extract” and Fig. 1B, “Extract, ×1/1”), indicating a loss of membrane fluidity. A moderate decrease in fluidity was observed in diluted extracts (Fig. 1B, ×1/30 and ×1/100, incubated for 40 min). However, longer incubation times led to a substantial decrease in fluorescence recovery (Fig. 1B, ×1/30 and ×1/100, incubated for 100 min). When heat-treated extracts were analyzed, the recovery curve showed a similar trend to the buffer control (Fig. 1C, “Extract, heated”). A bovine serum albumin (BSA) solution prepared to the same protein concentration in PB did not affect the fluorescence recovery (Fig. 1C, “BSA”). The protein concentration of 2.5 mg/mL in the extract was negligible compared to the background salt concentration, so the effect of osmotic pressure changes was probably negligible. The ability of cell extracts to alter membrane fluidity appears to be widespread among eukaryotes. A similar decrease in fluidity was observed with cytosolic extracts of human embryonic kidney cells (namely HEK293T), but at longer incubation times (Fig. 1D, “HEK293T”, incubated for 100 and 420 min). In contrast, no change in fluidity was observed with *Escherichia coli* cell extracts, even after 7 h of incubation (Fig. 1D, “*E. coli*” incubated for 420 min). The dependence on extract concentration and incubation time, as well as the thermolability, suggest that the loss of membrane fluidity was enzymatically induced.

**Figure 1.**
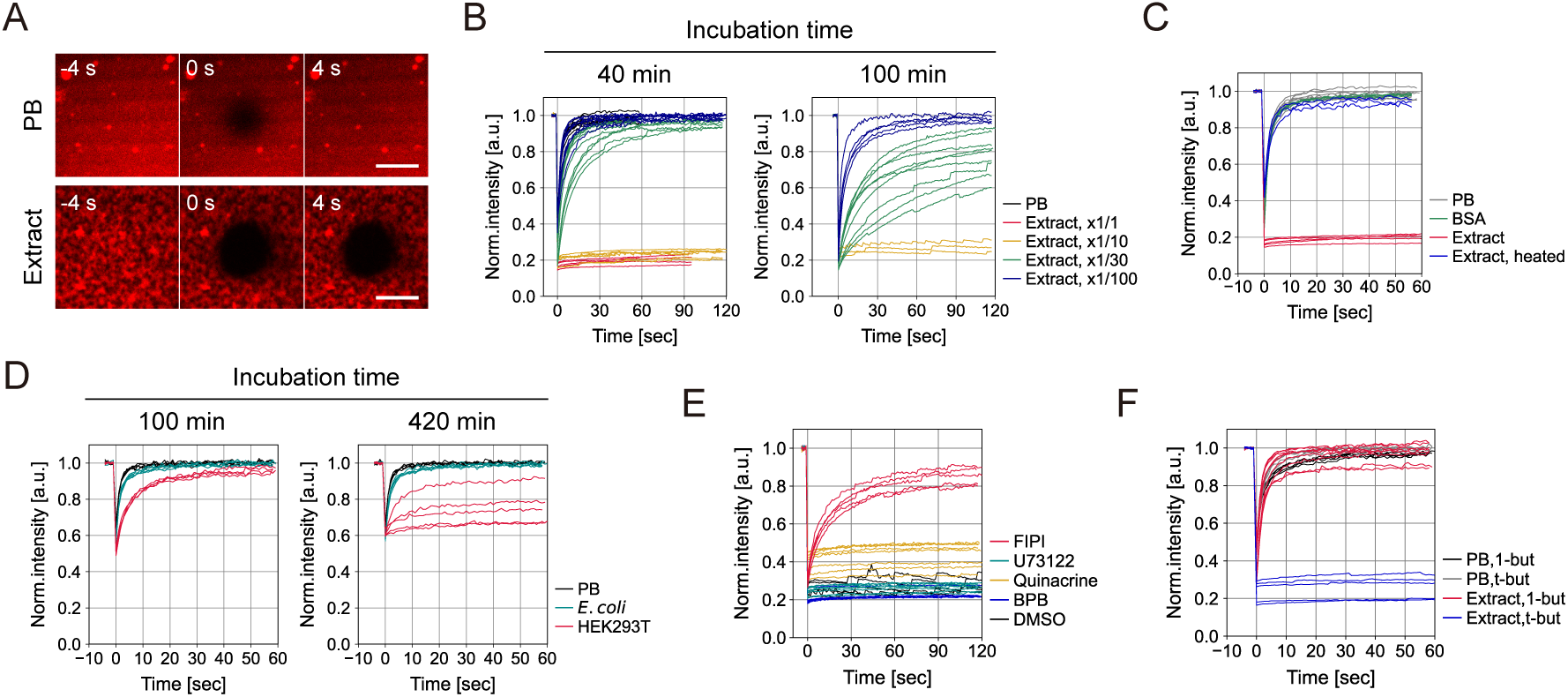
Phospholipase D present in cell extracts reduces lipid membrane fluidity. FRAP measurements of SLBs incubated with cell extracts. (**A**) Confocal images of SLBs composed of DOPC and 0.1% Rh-DMPE. Aqueous phases are phosphate buffer (PB) (*top*) and *Dictyostelium* AX2 cell extract (*bottom*). The central region (5 µm diameter) was photobleached at *t* = 0 s. Scale bars, 5 µm. (**B**) FRAP curves obtained for membranes incubated for 40 min (*left*) and 100 min (*right*) in PB and AX2 cell extracts diluted ×1/1–×1/100. (**C–F**) FRAP curves obtained for membranes in solution and cell extracts. (**C**) PB, 2.5 mg/mL BSA solution, AX2 cell extract, and heat-treated extract. Incubated for 40 min. (**D**) PB, *E. coli*, and HEK293T extracts. Incubated for 100 min (*left*) and 420 min (*right*). (**E**) AX2 extracts treated with phospholipase inhibitors: 500 µM FIPI, 100 µM U73122, 200 µM quinacrine, 300 µM BPB, and DMSO vehicle. (**F**) PB and AX2 extracts supplemented with 500 mM 1-butanol (1-but) and 500 mM t-butanol (t-but).

### PLD and its product phosphatidic acid (PA) induced loss of membrane fluidity

What causes the decrease in membrane fluidity? Because membrane fluidity is determined by lipid composition, the main candidate is phospholipases that hydrolyze phospholipids. We performed FRAP analysis of DOPC bilayers in cell extracts containing one of the following four phospholipase inhibitors: 5-fluoro-2-indolyl des-chlorohalopemide (FIPI), U73122, quinacrine or 4-bromophenacyl bromide (BPB). FIPI and U73122 target PLD and phospholipase C (PLC), respectively. Quinacrine and BPB are both phospholipase A2 (PLA2) inhibitors. Of these, only FIPI effectively blocked the loss of fluorescence recovery (Fig. 1E). FIPI treatment at concentrations above 300 µM maintained the mobile fraction for at least 5 h (Fig. S1B), during which the diffusion coefficient gradually decreased (Fig. S1A). PLD catalyzes the hydrolysis of PC to PA and choline. Because primary alcohols are stronger acceptors of phosphatidyl groups than water, the transphosphatidylation reaction favors the production of phosphatidylalcohols over PA. Addition of 1-butanol to the cytosolic extract prevented the loss of membrane fluidity (Fig. 1F, “Extract, 1-but”), whereas tertiary butanol did not (Fig. 1F, “Extract, t-but”). Mixing these alcohols with PB did not alter the recovery curves (Fig. 1F, “PB, 1-but” and “PB, t-but”). These results indicate that the generation of PA is essential for the loss of fluidity. Similar to PA, phosphatidylalcohols are generated by head group exchange. Therefore, the observed decrease in fluidity is not due to cleavage of the fluorophore from the lipid. This was further verified with another fluorescent lipid 18:1-12:0 nitrobenzoxadiazole (NBD)-PC. This lipid has a fluorescent group attached to one of the fatty acids, and using this probe, we also observed the reduced fluidity in cell extracts and its suppression by FIPI (Fig. S2).

### PA induced lipid condensation and slow diffusion depending on pH and Ca^2+^

PA contains a phosphomonoester headgroup with a unique structure that exhibits several properties, including a negative charge at physiological pH (pKa1 = 3.0, pKa2 = 8.0), hydrogen bonding, and a small headgroup that induces negative membrane curvature (24). Thus, accumulation of PA in SLBs is expected to increase the likelihood that membranes would detach from the substrate or rupture. We investigated the effect of PLD on DOPC membranes. In this minimal setup, the membranes became markedly heterogeneous over time (Fig. 2A). The sparse distribution of Rh-DMPE fluorescence was similar to that formed in cell extracts (comparison of Fig. 2A, “5 h” and Fig. 1A, “Extract” and Movie S1). Unlike the cytosolic extracts, numerous membranes detached from the glass surface and remained in the aqueous phase (Fig. 2B). This was likely due to incomplete lipid degradation. In membranes that remained attached to the glass surface, the diffusion coefficient and mobile fraction of Rh-DMPE gradually decreased, reaching a minimum at 5 h (Fig. 2C and D), recapitulating the loss of fluidity in cell extracts. The decrease in diffusion coefficient preceded that in mobile fraction (Fig. 2C and D, *t* = 1–2 h), suggesting that PA accumulation slows lipid diffusion before planar membrane disruption.

**Figure 2.**
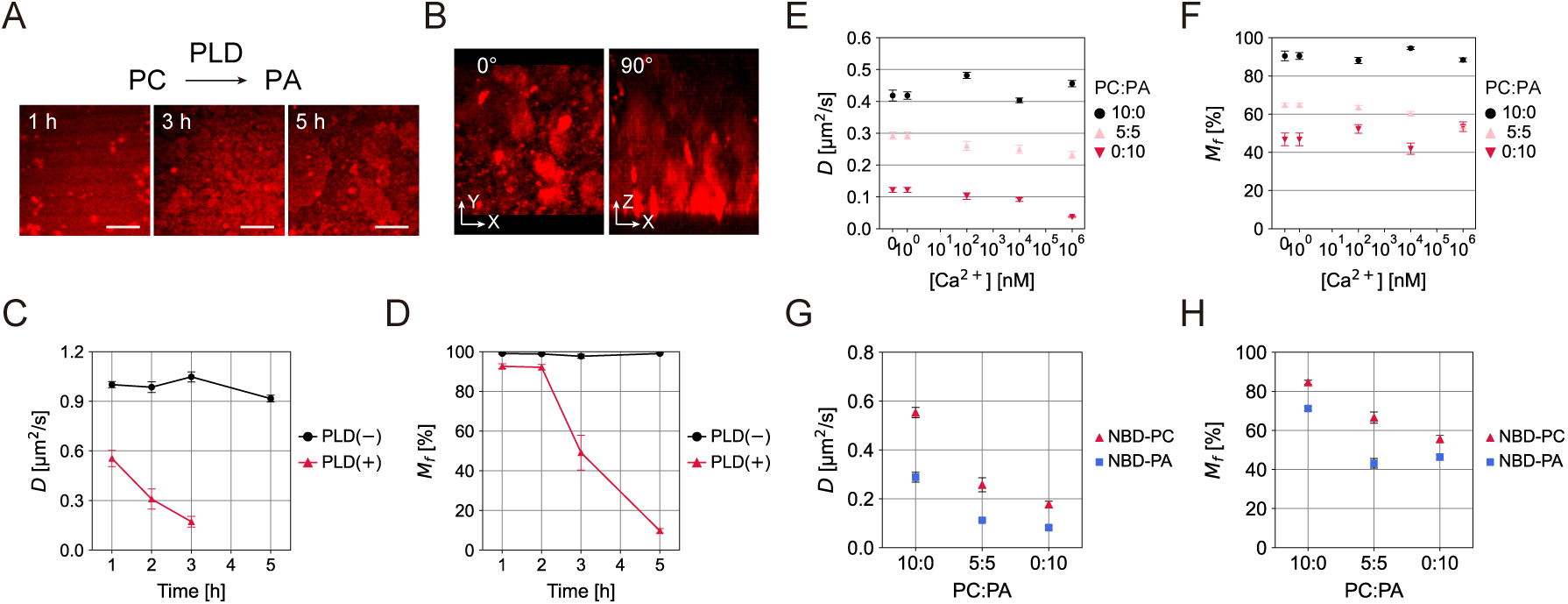
PA generated by PLD reduces lipid diffusivity. (**A** and **B**) Representative snapshots of Rh-DMPE in DOPC membranes during incubation with PLD. (**A**) Confocal images taken close to the glass surface. Numbers, incubation times. Scale bars, 5 µm. (**B**) Maximum intensity projections of Zstacks acquired after 5 h of incubation. Projections into the XY (*left*) and XZ (*right*) planes. (**C** and **D**) *D* (**C**) and *M_f_* (**D**) of Rh-DMPE (mean ± s.e., N = 5–10 per condition). (**E–H**) FRAP measurements of DOPC and DOPA films on PLL-coated coverslips. (**E** and **F**) Ca^2+^ dependence of Rh-DMPE. (**E**) *D* and (**F**) *M_f_* (mean ± s.e., N = 10–20 per condition). Legend, molar ratio PC:PA. (**G** and **H**) PC:PA ratio dependence of NBD-PC and NBD-PA. (**G**) *D* and (**H**) *M_f_* (mean ± s.e., N = 5–15 per condition).

The effect of PA on membrane fluidity was evaluated by preparing bicomponent lipid films consisting of DOPC and DOPA. Divalent cations counteract the electrostatic repulsion between anionic phospholipids, thereby affecting the lateral diffusion of lipids (25–27). However, conventional planar membrane formation methods require the addition of millimolar Ca^2+^ to facilitate the adsorption of lipid vesicles to glass surfaces. Therefore, we developed a Ca^2+^-free method using glass surfaces coated with the cationic polymer poly-L-lysine (PLL) (Fig. S3A) (see *SI Appendix*, *Materials and Methods* for details). Unlike the DOPC vesicles that adsorbed and ruptured on the PLL-coated surface, DOPA vesicles remained condensed (Fig. S3B). Therefore, we obtained planar films by increasing the pH to 10.0 to fully deprotonate PA (Fig. S3C) and replacing the aqueous phase with PB (pH 6.5) containing defined Ca^2+^ concentrations before FRAP measurements (Fig. 2E and F). Notably, the mobile fraction of Rh-DMPE was nearly 100% in PC (Fig. 2F, “10:0”), but decreased to about 50% in PA membranes (Fig. 2F, “5:5” and “0:10”). This suggests that PA in the lower leaflet strongly interacts with PLL and is immobilized, whereas lipids in the upper leaflet are freely diffusing. The measured diffusion coefficients were consistently lower in PA membranes than in PC membranes (Fig. 2E). Lipid diffusivity in PA membranes was reduced by high Ca^2+^ concentrations, considerably in the millimolar range (Fig. 2E). In contrast, no Ca^2+^-induced decrease in diffusivity was observed in PC membranes (Fig. 2E). To directly compare lipid diffusion with different head groups, we used two fluorescent lipids, 18:1-12:0 NBD PC (NBD-PC) and 18:1-12:0 NBD PA (NBD-PA) (Fig. 2G and H). Both lipid probes were added to lipid films at 0.1 mol%, and diffusion slowed as the PA ratio increased (Fig. 2G). The diffusivity of NBD-PA was consistently lower than that of NBD-PC, indicating that PA itself diffused slowly within the constructed membranes.

### PLD altered the diffusivity of a myristoylated domain of PKBR1 in living cells

The *in vitro* experiments described above demonstrated a slowdown in lipid diffusion in PA-enriched membranes. We next investigated whether PLDs alter protein diffusion in membranes of living cells. *Dictyostelium* PldB is a mammalian PLD1/2 homolog (Fig. 3A) (28, 29) and is present in the plasma membrane (Fig. 3B, “PldB_FL_”) (30). Based on the domain organization (Fig. 3A), we expressed five types of truncated PldB with an N-terminal mScarlet-I tag (Fig. 3B) and examined their association with the plasma membrane. The PH domain (346–464 aa) did not localize to the plasma membrane (Fig. 3B, “PldB_PH_”), and the large N-terminal fragment (1–464 aa) containing the calcium-binding EF-hand-like motifs showed weak binding (Fig. 3B, “PldB_N1–464_”). The shorter N-terminal fragment (1–346 aa) was distributed mainly in the cytoplasm (Fig. 3B, “PldB_N1–346_”), suggesting that the EF-hand region and the PH domain function together for membrane binding. Fragments containing the putative phosphatidylinositol 4,5-bisphosphate (PIP2)-binding region (624–721 aa) or the C-terminal catalytic domain (886–1216 aa) were found in the cytoplasm (Fig. 3B, “PldB_PIP2_” and “PldB_cat_”). These observations indicate that *D. discoideum* PldB binds to the plasma membrane through various domains, similar to mammalian PLD1/2, which binds to the membrane via palmitoylation (31) and electrostatic interactions (32). Note that as PldB is not responsible for the loss of SLB fluidity in cytosolic extracts, since it is mainly present in the plasma membrane. Extracts from *pldB*^−^ cells showed similar effects to those from wild-type cells (Fig. S4). To examine the effects of PLD on the plasma membrane, we focused on PldB.

**Figure 3.**
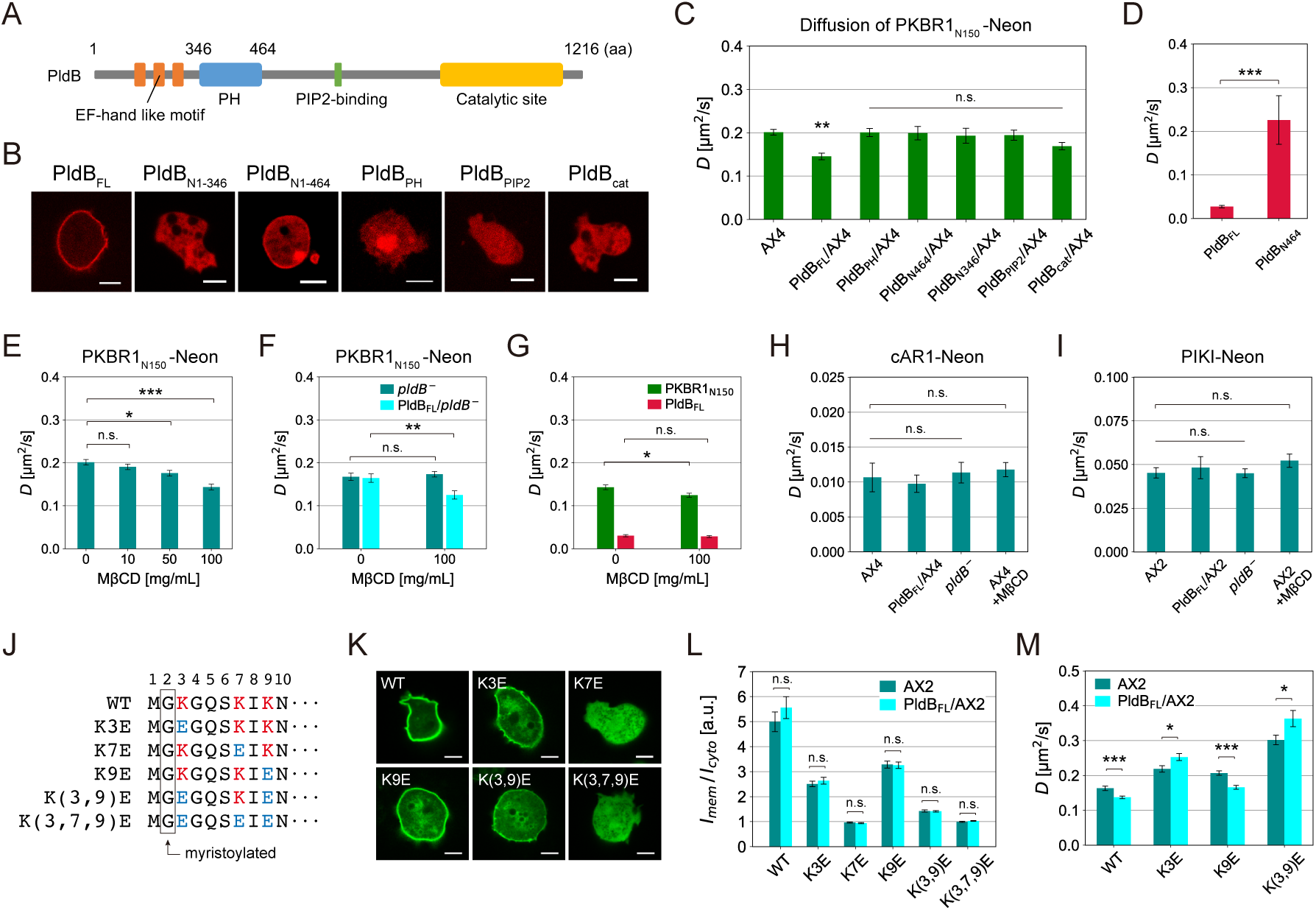
PLD reduces the diffusivity of the myristoylated domain of PKBR1 in living cells. (**A**) Domain organization of PldB. (**B**) Confocal images at the equatorial plane of AX4 cells expressing PldB and its truncated versions tagged with mScarlet-I (mSca) at the N-terminus. Subscripts, FL: Full-length, PH: 346–464 aa, PIP2: 624–721 aa, cat: 886–1216 aa. Scale bars, 5 µm. (**C–I**) *D* of membrane proteins measured by FRAP (mean ± s.e.). (**C**) PKBR1_N150_-Neon in AX4 and cells overexpressing the PldB constructs shown in **B** (N = 18, 27, 14, 17, 18, 19, and 20 cells). (**D**) mSca-PldB_FL_ and mSca-PldB_N1–464_ in AX4 (N = 17 and 15 cells). (**E**) PKBR1_N150_-Neon in AX2 treated with MβCD (N = 18, 20, 21, and 14 cells). (**F**) PKBR1_N150_-Neon in *pldB*^−^ and cells expressing mSca-PldB_FL_ (“PldB_FL_/*pldB*^−^”). Cells were treated with MβCD (N = 30, 16, 29, and 20 cells). (**G**) PKBR1_N150_-Neon and mSca-PldB_FL_ in AX4 overexpressing mSca-PldB_FL_. Cells were treated with MβCD (N = 25, 16, 22, and 21 cells). (**H**) cAR1-Neon in AX4, AX4 overexpressing mSca-PldB_FL_ (“PldB_FL_/AX4”), *pldB*^−^ and AX4 treated with 100 mg/mL MβCD (N = 12, 13, 9, and 15 cells). (**I**) PIKI-Neon in AX2, AX2 overexpressing mSca-PldB_FL_ (“PldB_FL_/AX2”), *pldB*^−^ and AX2 treated with 100 mg/mL MβCD (N = 23, 19, 24, and 15 cells). (**J**) N-terminal sequences of original PKBR1 (“WT”) and mutated constructs. (**K**) Confocal images at the equatorial plane of AX2 cells expressing PKBR1_N150_-Neon or its mutants. Scale bars, 5 µm. (**L**) Membrane-to-cytosol intensity ratios of the Neon-tagged fragments shown in **K** (mean ± s.e., N = 10, 10, 12, 8, 12, and 10 cells for AX2 and 10, 14, 12, 14, 15, and 15 cells for PldB_FL_/AX2). (**M**) *D* of myristoylated PKBR1 fragments expressed in AX2 (N = 19, 18, 18, and 19 cells) and PldB_FL_/AX2 (N = 24, 20, 19, and 16 cells). Data were compared using Student’s t-test for two groups and one-way ANOVA followed by Kramer–Tukey’s post-hoc test for more than two groups. \**p* < 0.05, \*\**p* < 0.01, and \*\*\**p* < 0.001. n.s., not significant.

To measure the lateral diffusion of membrane proteins in live cells, we used *Dictyostelium* cells expressing mNeonGreen (Neon) fused to an N-terminal fragment of AKT/SGK homolog PKBR1 (33), which is anchored to the inner leaflet of the plasma membrane via N-myristoylation (Fig. S5A) (34). FRAP analysis showed that the diffusion coefficient of PKBR1_N150_-Neon in wild-type cells was 0.201 ± 0.007 µm^2^/s (Fig. 3C, “AX4”). Treatment with sodium azide, which nonspecifically inhibits ATP-dependent processes including endocytosis, had no significant effect (Fig. S5B). This indicates that the lateral diffusion of PKBR1_N150_-Neon was sufficiently fast compared to its dissociation rate from the plasma membrane. This is consistent with the reported mean residence time of about 40 s (3). In cells overexpressing full-length PldB, we found that the diffusion coefficient of PKBR1_N150_-Neon was significantly reduced to 0.15 ± 0.01 µm^2^/s (Fig. 3C, “PldB_FL_/AX4”). This reduction was not observed in cells expressing PldB fragments (Fig. 3C). The diffusion coefficient of mScarlet-I-PldB_FL_ was 0.026 ± 0.003 µm^2^/s (Fig. 3D), approximately 8 times slower than that of PKBR1_N150_-Neon. In comparison, the diffusion coefficient of mScarlet-I-PldB_N464_ was 0.23 ± 0.06 µm^2^/s, almost an order of magnitude higher (Fig. 3D). Taking into account the weak membrane localization (Fig. 3B, “PldB_N1–464_”), this is an overestimate since it does not consider the presence of reversible membrane binding. Overall, lateral diffusion of PKBR1_N150_ was reduced in cells with elevated PldB levels, for which the C-terminal catalytic domain of PldB is required.

It has been recently reported that polyphosphates contained in extracellular secretions reduces membrane fluidity (35). To examine cell secretions, the supernatant of the cell suspension was collected after 1 h of starvation and used as conditioned medium (CM). After incubation in CM for 30 min, the diffusivity of PKBR1_N150_-Neon was significantly decreased (Fig. S6A, “CM(+)”), indicating that CM affects protein diffusion on the inner membrane. Because heat-treated CM was less effective in decreasing diffusion (Fig. S6A, “CM, heated(+)”), CM may contain factors other than polyphosphates that reduce membrane fluidity. The diffusivity of PKBR1_N150_-Neon in *pldB*^−^ was not significantly decreased by CM (Fig. S6B). These results indicate that PldB is required for the reduction of PKBR1 diffusion in CM.

### PldB selectively reduced diffusion of the myristoylated domain of PKBR1 in a sterol-dependent manner

Metazoan PLDs are commonly found in cholesterol-rich membrane rafts and their activity increases when rafts are disassembled (36, 37). *D. discoideum* does not synthesize cholesterol but does produce stigmasterol (38, 39), whose function is poorly understood except that it is known as the precursor of the sporulation-inducing steroid SDF-3 (40). Genes required for sterol synthesis, *cas1* and *smtA*, encoding cycloartenol synthase and sterol methyltransferase, respectively, are transiently increased in expression 3–5 h after development (Fig. S7A and B). To deplete sterols, we used methyl-β-cyclodextrin (MβCD), a cyclic heptamer of D-glucose that strongly interacts with sterols in its hydrophobic cavity and is widely used to deplete cholesterol and phytosterols (41). On agar plates containing MβCD, the onset of cell aggregation was delayed in a concentration-dependent manner (Fig. S7C). While it took 6 h for cells on plain agar to begin streaming (Fig. S7C, “0 mg/mL”), the timing of this event was delayed by 4 h on 100 mg/mL MβCD-containing agar (Fig. S7C, “100 mg/mL”). Since cell streaming itself appeared normal on MβCD-containing agars, early developmental timing was mostly affected, rather than cell migration or cell–cell adhesion. Moreover, at 33 mg/mL, fruiting bodies showed aberrant morphologies with short stalks (Fig. S7C). At 100 mg/mL, culmination was completely arrested (Fig. S7C).

At the single-cell level, MβCD treatment markedly reduced the diffusion of PKBR1_N150_-Neon (Fig. 3E). No such effect was observed in *pldB*^−^ (Fig. 3F, “*pldB*^−^”), and this effect was reverted by PldB overexpression in the null background (Fig. 3F, “PldB_FL_/*pldB*^−^”). These data indicate that MβCD acts on PldB to reduce the diffusivity of PKBR1_N150_. The effects appeared to be additive, since the reduction in the diffusivity of PKBR1_N150_-Neon in PldB overexpressors (Fig. 3C) was further accentuated by MβCD treatment (Fig. 3G, “PKBR1_N150_”). In contrast, MβCD treatment did not alter the diffusivity of mScarlet-I-PldB_FL_ (Fig. 3G, “PldB_FL_”). To test the generality, two other membrane proteins were studied. The diffusion coefficient of Neon-fused G protein-coupled receptor cAR1 (Fig. S5C) was 0.011 ± 0.002 µm^2^/s (Fig. 3H, “AX4”), which is consistent with the value measured by SPT (42). The diffusion rate of cAR1-Neon was independent of PldB and sterols, since it was not altered by overexpression or knockout of PldB (Fig. 3H, “PldB_FL_/AX4” and “*pldB*^−^”) and MβCD treatment (Fig. 3H, “AX4+MβCD”). Phosphatidylinositol-4-phosphate 5-kinase (PI4P5K) PIKI tagged with Neon was localized on the plasma membrane (Fig. S5D), and its diffusion coefficient of 0.045 ± 0.003 µm^2^/s was also not significantly altered (Fig. 3I). The proteins Gα2 and Arf could not be tested because their low expression levels did not allow a reliable fluorescence readout. Nevertheless, these results suggest that the changes in diffusivity caused by PldB and sterol depletion are not generic for all membrane proteins.

Myristoylated proteins such as Src (43) and MARCKS (44), require a polybasic region adjacent to the lipidated end for sufficient membrane targeting. Similarly, the N-terminus of PKBR1 contains three lysine residues within the first 10 amino acids. To test their contribution to membrane binding and lateral diffusion, we systematically replaced them with glutamic acid (Fig. 3J). Compared with unmutated PKBR1_N150_-Neon (Fig. 3K and L, “WT”), single substitutions of either K3E or K9E reduced the membrane-to-cytosol fluorescence ratio (Fig. 3K and L, “K3E” and “K9E”). Double substitutions further weakened membrane binding (Fig. 3K and L, “K(3,9)E”). In contrast, the K7E mutation completely abolished membrane localization (Fig. 3K and L, “K7E” and “K(3,7,9)E”), suggesting its requirement for N-myristoylation. Co-expression of PldB_FL_ did not significantly affect the membrane-to-cytosol ratio of these mutant fragments (Fig. 3L, “AX2” and “PldB_FL_/AX2”), suggesting that their localization to the plasma membrane is not strongly dependent on PA. We then compared the diffusion rates of these mutant fragments between parental AX2 and PldB overexpressors (Fig. 3M, “AX2” and “PldB_FL_/AX2”). In wild-type AX2, the diffusion rate increased with the number of mutations (Fig. 3M, “AX2”), showing an inverse trend with the the membrane-to-cytosol ratio, likely due to enhanced dissociation from the membrane. In the K9E mutant, a decrease in the diffusion rate with PldB overexpression was observed (Fig. 3M, “K9E”). However, in the K3E-substituted fragments, the diffusion rates increased with PldB overexpression (Fig. 3M, “K3E” and “K(3,9)E”). This suggests that repulsive electrostatic interactions at the protein– membrane interface accelerate lateral diffusion. These results indicate that lysine residues additionally contribute to the interaction of the N-terminus of PKBR1 with anionic phospholipids for membrane targeting, in which the proximal lysine serves critically to reduce the diffusion rate in PA-enriched membranes.

### PldB regulates PI(3,4,5)P3 amplification in macropinocytic patches and chemotactic responses

Given its specific effect on PKBR1 diffusion, we addressed whether PldB controls downstream signaling of PKBR1 (Fig. 4A). PI4P5K (PIKI) is a major effector of PKBR1 (45). GPCR-mediated activation of mTORC2 and PDK1 activates PIKI to generate PI(4,5)P2 (45–48). The basal level of PI(4,5)P2 is critical for the amplification of PI(3,4,5)P3 (13, 48, 49). Therefore, we anticipated that PldB would participate in PI(3,4,5)P3 signaling. PI(3,4,5)P3 forms micron-sized patches in the membrane through an excitatory signaling network that controls the activity of Ras and PI3K (13, 15, 16). We used PH_CRAC_-GFP to monitor the PI(3,4,5)P3 patches formed on the ventral side of the cells (Fig. 4B). Before patch formation, the membrane was uniformly covered with mScarlet-I-PldB (Fig. 4B, t = 0 s). As the PH_CRAC_-GFP-enriched patch expanded, mScarlet-I-PldB was depleted (Fig. 4B, t = 60 s). In PldB-overexpressing cells, the frequency of patch occurrence was reduced (Fig. 4C and D, “AX2” and “PldB/AX2”). In *pldB*^−^, patch occurrence was increased (Fig. 4C and D, “*pldB*^−^”), which was suppressed by PldB overexpression (Fig. 4C and D, “PldB/*pldB*^−^”). The size and duration of the patch were consistently reduced with PldB overexpression (Fig. S8). These results indicate that PldB inhibits PI(3,4,5)P3 patches.

**Figure 4.**
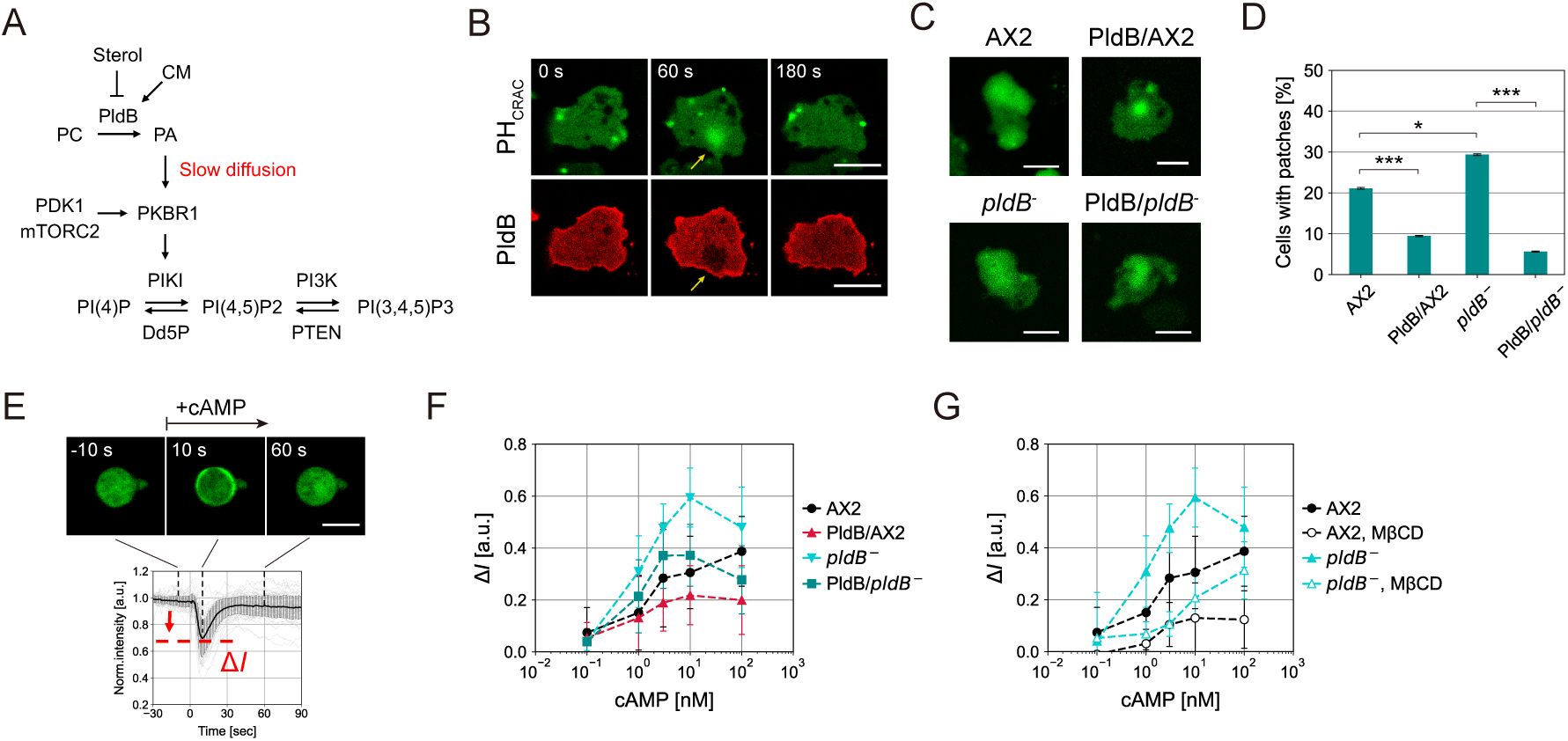
PLD controls PIP3 amplification in propagating patches and the chemotactic responses. (**A**) PKBR1 signaling. (**B**) Confocal images of PH_CRAC_-GFP (*top*) and mScarlet-I (mSca)-PldB (*bottom*) taken at the ventral surface. Yellow arrows indicate a PI(3,4,5)P3 patch. (**C**) Confocal images of PH_CRAC_-GFP in AX2, AX2 overexpressing mSca-PldB (“PldB/AX2”), *pldB*^−^ and *pldB*^−^ expressing mSca-PldB (“PldB/*pldB*^−^”). Scale bars, 10 µm. (**D**) Percentage of cells with PH_CRAC_-GFP patches (mean ± s.e., N = 139, 116, 139, and 110 cells). Data were compared using one-way ANOVA followed by Kramer–Tukey’s post-hoc test. \**p* < 0.05 and \*\*\**p* < 0.001. (**E**) Confocal images of PH_CRAC_-GFP (*top*). cAMP was added uniformly at *t* = 0 s. Temporal profile of PH_CRAC_-GFP intensity at cytosol (*bottom*). (**F** and **G**) Intensity drop of cytosolic PH_CRAC_-GFP (Δ*I*) upon cAMP stimulation at various concentrations (mean ± s.d.). Cells were pretreated with 5 µM Latrunculin A. (**F**) Dose-response curves for AX2, PldB/AX2, *pldB*^−^, and PldB/*pldB*^−^ cells. (**G**) Dose-response curves for AX2 and *pldB*^−^ treated with 0 and 100 mg/mL MβCD.

PI(3,4,5)P3 is transiently produced when cells are stimulated with cAMP, inducing the translocation of PH_CRAC_-GFP from the cytosol to the plasma membrane (Fig. 4E). We measured the intensity of PH_CRAC_-GFP during cAMP stimulation (Fig. S9), from which we quantified the average intensity decrease (Δ*I*) in the cytosolic region for each strain. Cells were pretreated with Latrunculin A to disrupt the actin cytoskeleton. Overexpression of PldB suppressed the membrane translocation of PH_CRAC_-GFP after cAMP stimulation (Fig. 4F, “AX2” and “PldB/AX2”), whereas it was potentiated in *pldB*^−^ (Fig. 4F, “*pldB*^−^”). Expression of PldB in *pldB*^−^ reduced the response to a level comparable to that of the parental wild-type (Fig. 4F, “PldB/*pldB*^−^”). We also found that MβCD treatment inhibited membrane translocation of PH_CRAC_-GFP in response to cAMP stimulation (Fig. 4F, “AX2” and “AX2, MβCD”). Interestingly, MβCD treatment severely inhibited the response in *pldB*^−^ (Fig. 4G, “*pldB*^−^” and “*pldB*^−^, MβCD”), suggesting that sterol depletion has a different effect independent of PldB. Overall, these data consistently indicate that PldB inhibits PI(3,4,5)P3 production in the membranes.

## Discussion

In this study, we demonstrated that cytosolic extracts triggered a marked decrease in the fluidity of SLBs and that PLD and its product PA were responsible for this phenomenon. The phosphate headgroup of PA forms hydrogen bonds with the surrounding lipids, increasing the lipid packing density (50, 51). Because PA is conical, PA-rich regions should naturally exhibit curvature. Therefore, PA accumulation may generate lipid-condensed regions that easily detach from the solid support, disrupting the planar membrane. To directly measure lipid diffusion in PA-rich membranes, we prepared supported films on PLL-coated substrates. The lateral diffusion of lipids was markedly reduced at high PA ratios, and the diffusion of PA itself was slow. Since the cell extracts used in this study were collected from the cytosol, the influence of lipid modulators located on the membrane, including PldB, was not responsible for the slowdown observed in the model membrane. Moreover, the PLD inhibitor only partially blocked the reduction in lipid diffusivity (Fig. S1A). This suggests the presence of other factors that may alter diffusion, such as other phospholipases and desaturases. Our observations may also be related to the bacteriolytic activity of cell extracts, for which putative antibacterial effectors have been detected using proteomic analysis (52).

Studies of model membranes have shown that there is lateral phase separation between PC and PA in the presence of divalent cations (25–27, 53, 54). However, intracellular PA concentrations are relatively low (less than 1% of the total phospholipids) (23), and it is unclear whether large PA-rich domains can exist in living cells. FRAP measurements show spatially averaged viscosity over micrometer lengths, and therefore do not indicate whether the PA forms raft-like compartments or is homogeneously scattered. This is also due to the lack of noninvasive probes for PA, which shares the same problem as conventional lipid rafts. Although the structural basis remains unclear, our results demonstrate that PLD reduces the diffusivity of the myristoylated fragment of PKBR1. Among the three proteins tested, PLD-induced slowing was observed only for PKBR1, as confirmed by PLD overexpression and sterol depletion. Such selectivity suggests that the direct interaction with PA, rather than increased packing density of lipids, underlies the reduced diffusivity. As the cationic polymer poly-lysine could induce lipid phase separation by crosslinking PA (26), it is possible that the polybasic region near the myristoylated tail is responsible for the interaction with PA. Our findings indicate that residues 3K and 9K of PKBR1 contribute to membrane binding, supporting multipoint interactions with anionic phospholipids.

Membrane binding through a combination of lipidation and electrostatic interactions may allow dynamic control of membrane affinity (55–58). The “myristoyl-electrostatic switch” mechanism refers to the translocation of proteins to the cytosol upon phosphorylation of serine or threonine residues near the lipidated end. Findings from this study suggest that a similar effect occurs laterally, modulating lateral diffusion depending on surface charge (Fig. 3L and M). This likely underlies the macroscopic segregation on membranes. According to a recent SPT study, PKBR1 exhibits bimodal lateral diffusion on the plasma membrane (3). When a PI(3,4,5)P3-rich patch expands, PKBR1 molecules within the patch diffuse mostly in a fast mode, facilitating the translocation of this kinase out of the patch (3). The outer region of the patch is characterized by a relatively high negative surface charge (17) and enrichment of PldB, which likely contributes to providing a surface charge for capturing PKBR1. The same mechanism may also act for PldB that is partitioned outside the patches if PldB binds to membranes by palmitoylation (31) and electrostatic interactions (32), similar to mammalian PLD1/2. Moreover, membrane curvature strongly influences partitioning dynamics: in cells interfaced with micrometer-scale ridges, PI3,4,5)P3-rich patches are localized around the ridge structures (16). This may be explained by PA preferring to accumulate in positively curved regions, facilitating its depletion from the surface attached to the ridges.

The results of this study suggest that PldB-induced slow diffusion of PKBR1 not only serves for its partitioning but also regulates phosphoinositide signaling. A study of alcohol-exposed cells suggested that PLD positively regulates PI(4,5)P2 (59). PKBR1 activates PIKI (45) and its product PI(4,5)P2 is required for PKBR1 phosphorylation, likely constituting a positive feedback loop (48). PI(4,5)P2 content is crucial for PI(3,4,5)P3 amplification (13, 48, 49). PLC controls PI(4,5)P2 levels by hydrolysis, increasing PI(3,4,5)P3 produced in response to cAMP stimuli (49). Moreover, synthetically induced PI(4,5)P2 reduction lowers the threshold for PI(3,4,5)P3 production (13). In this study, we demonstrated that PldB suppresses the occurrence, size, and lifetime of PI(3,4,5)P3 patches, as well as the maximum amount of PI(3,4,5)P3 produced in response to cAMP. These results suggest that PldB-induced slow diffusion of PKBR1 increases the basal level of PI(4,5)P2 and antagonizes the transient production of PI(3,4,5)P3. The enhanced activity in slow kinetics suggests the complexation of PKBR1 with lipids or other proteins. In this regard, our results could not indicate whether PLD-induced slow diffusion is due to an increase in the population of slower modes or an overall shift toward the slower. Implications may be gained from further SPT studies.

The reduced diffusivity of PKBR1 in sterol-depleted cells suggests that PldB activity depends on sterols. Sterols play regulatory and structural roles in lateral organization of membranes (60, 61). In metazoan cells, palmitoylated PLD is inactivated by sequestering it from its substrate PC by cholesterol (36, 37). *Dictyostelium* cells grow as single cells and aggregate upon starvation to form multicellular structures. The transition from the growth stage to the onset of aggregation takes several hours, during which the cells differentiate and secrete molecules (62). As *pldB*^−^ cells develop precociously (28, 29), PldB is thought to set a threshold for aggregation. An increase in the amount of sterols during the aggregation phase (38) (Fig. S7A and B) may suppress PLD activity, thereby enhancing the cellular sensitivity to cAMP. MβCD treatment severely delayed the onset of aggregation and inhibited the response to cAMP, though we might be careful in the specificity of this treatment (63). Despite repeated attempts, we were unsuccessful in creating a *cas1* knockout strain, suggesting that sterols are essential for growth. Together with recent findings that Ras activation is suppressed in membranes with excess sphingomyelin (64), sterols and PLDs appear to constitute the key regulators of reaction and diffusion dynamics in membranes.

## Materials and Methods

All the plasmids and cell strains used in this study, along with experimental procedures for preparing cell extracts and lipid films, live-cell imaging, FRAP measurements, and image analyses are described in the *SI Appendix*, *Materials and Methods*.

## Author Contributions

GH conceived, planned, and managed all aspects of the project. GH performed all the experiments and analyses. GH and CT performed fluorescence recovery after photobleaching measurements on lipid films. GH, SS, and MY wrote the manuscript.

## Classification

Biological Sciences/Biophysics.

## Acknowledgments

We thank Prof. Taro QP Uyeda and the National BioResource Project (NBRP) Nenkin for their kind gift of *pldB*^−^. We thank Hiroyasu Kamei for kindly providing HEK293T cells. We thank Hidenori Hashimura for technical support with plasmid construction. We thank Nao Shimada for technical support in preparing the cell extracts. This work was supported by grants from the Japan Science and Technology Agency (JST), CREST (JPMJCR1923, S. S.), FOREST (JPMJFR213Y, M. Y.), and the Japan Society for the Promotion of Science (JSPS) KAKENHI (JP22H01188, M. Y. and JP19H05801, S. S.).

## Data Availability

All data included in this study will become available from the RIKEN Systems Science Biological Dynamics database repository.

