## Supplementary text for "Slow diffusion and signal amplification on membranes regulated by phospholipase D"

### Slow diffusion and signal amplification on the membrane regulated by phospholipase D

**This PDF file includes:**

Supporting text
Table S1 to S2
Figures S1 to S9
Legend for Movie S1
SI References

**Other supplementary materials for this manuscript include the following:**

Movie S1

#### Supporting text

#### Materials and Methods

##### Plasmids and cell strains

Plasmid pDM358 containing the Dicty-codon-optimized mScarlet-I, pDM358 mScarlet-I-MCS, was a kind gift from Dr. Hashimura (NBRP Nenkin, G90683). The PldB open reading frame was cloned into this following the in-fusion kit protocol (Takara, 638948). Fragments of PldB, 1-346 aa, 1-464 aa, PH domain (347-464 aa), 625-720 aa and 887-1216 aa, were cloned into pDM358 mScarlet-I-MCS. These gene fragments were expressed under the *act15* promoter, except when expressed under the *coaA* promoter in Fig. 3L-M. cAR1 and PIKI were cloned into plasmid pDM304 MCS-mNeonGreen (Neon) and expressed under the *act15* promoter. The N-terminal fragment of PKBR1 (1-150 aa) was cloned into pDM304 MCS-Neon and expressed under the *coaA* promoter. The mutant strain *pldB*$^-$ was a kind gift from Prof. Taro Uyeda (NBRP Nenkin, S00559) (1). Laboratory wild-type strains AX4 and AX2, and the mutant strain *pldB*$^-$ were transformed by the standard electroporation method. The plasmids and cell strains used in this study are listed in Table. S1-2.

##### Preparation of cell extracts

*Dictyostelium discoideum* cells were cultured as previously described (2). To collect cell extracts, $1\times10^8$ AX2 cells were washed twice and resuspended in 1 mL phosphate buffer (PB). Using a cell disruption vessel (Model 46391, Parr Instrument), cells were pressurized with nitrogen to 350 psi and left on ice for 15 min. The extruded cell lysate was centrifuged twice for 30 min at 16,000 rcf, and the cytosolic fraction was obtained from the supernatant. HEK293T cells were cultured in D-MEM (Nissui, 05915) supplemented with 10% FBS and 0.45% glucose at 37℃ and 5% $CO_2$. $3\times10^7$ cells were collected and washed with PBS buffer, and the cytosolic fraction was obtained by ultrasonic cell disruption (Tomy, UD-201) and centrifugation. *Escherichia coli* B/r cells were cultured overnight in 20 mL of LB medium. The cells were washed and resuspended in 1 mL of PB. The cytosolic fraction was obtained by ultrasonic cell disruption and centrifugation. The total protein concentration in cell extracts was measured by Bradford protein assay (Biorad, 500-0006JA) and adjusted to 2.5 mg/mL with buffer. The concentration of phospholipids remaining in the AX2 extracts was measured using a kit (Fujifilm Wako, 296-63801), which was below the detection limit of 4 μg/mL. Extracts were stored at 4℃ and used within 3 days.

Phospholipase inhibitors FIPI (Calbiochem, 528245), U73122 (Cayman Chemical, 70740), quinacrine (Abcam, ab120749) and 4-bromophenacyl bromide (BPB; TCI, A5501) were used for drug treatment of extracts. FIPI, U73122 and BPB were dissolved in DMSO at 10 mM, 1 mM and 30 mM, respectively, and quinacrine was dissolved in water at 10 mM. Aliquots were stored at −20℃. 1-butanol and tert-butanol (Nacalai tesque, 06015-95 and 06103-35) were stored at room temperature. Cell extracts were treated with these reagents for 30 min at 22℃ before application to supported lipid bilayers. Heat treatment of the cell extract was carried out at 80℃ for 30 min.

**Preparation of lipid films**

1,2-dioleoyl-sn-glycero-3-phosphatidylcholine (DOPC) and 1,2-dioleoyl-sn-glycero-3-phosphate (DOPA) lipids (Avanti polar lipid, 850375P and 840875P, $T_m$ = −17℃ and −4℃) were dissolved in chloroform at a concentration of 10 mg/mL and stored at −20℃. The fluorescent lipids, 14:0 Liss Rhod PE (Rh-DMPE; Avanti, 810157P), 18:1-12:0 NBD PC (Avanti, 810133P) and 18:1-12:0 NBD PA (Avanti, 810176P) were dissolved in chloroform at 1 mg/mL and stored at −20℃. 0.64 μmol of lipids were mixed in a tube with 0.1 mol% fluorescent lipid, and the solvent was evaporated under air flow to form a lipid film. This was resuspended in 1 mL of ultrapure water and sonicated in an ice-water bath for 5 min to obtain the SUV solution. The solution was stored at −20℃. Coverslips washed with ethanol and NaOH were treated with air plasma (Harrick Plasma, PDC-32G) to increase hydrophilicity, and then covered with SUVs solution supplemented with 5 mM $CaCl_2$. After incubation at room temperature for 1 h, the substrate was washed thrice with PB (pH 6.5).

To prepare $Ca^{2+}$-free lipid films, glass-bottom dishes (Mattek, P35G-0-14-C) were treated with air plasma and covered with 100 μg/mL poly-L-lysine (PLL) solution (Sigma, P4832). After 1 h of incubation, the surface was washed with copious amounts of water and dried. The surface was then covered with SUVs solution for 15 min, after which 100 mM $NaHCO_3$-NaOH buffer was added to adjust the pH to 10.0 to fully deprotonate the PA lipids. After 1 h of incubation, the surface was washed thrice with PB. The specified concentrations of $CaCl_2$ aq were added at least 30 min before measurements.

Lipid films formed on coverslips were incubated with cell extracts, buffer and BSA solution at 22℃ for 40 min (or longer times as indicated in figure legends) prior to FRAP measurements. For extracts from *E. coli* and HEK293T cells, membranes were incubated at 37℃ for 7 h. Phospholipase D (PLD) from *Arachis hypogaea* (Sigma, P0515) was dissolved in phosphate buffer at 100 mg/mL and stored at −20℃. 10 mg/mL PLD solution was applied to the membrane, and FRAP measurements were performed during incubation at 37℃.

**Live-cell imaging**

For live-cell imaging and FRAP measurements, cells were washed twice and resuspended in development buffer (DB) at $1\times10^7$ cells/mL. After shaking at 22℃, 155 rpm for 1 h, cells were plated on glass-bottom dishes (Mattek, P35G-0-14-C) at approximately $2\times10^4$

cells/cm$^2$. Methyl-β-cyclodextrin (MβCD) was purchased from Fujifilm Wako (320-84252). Conditioned medium (CM) was collected from the supernatant of cell suspensions shaken in DB for 1 h. Heat treatment of CM was performed at 95℃ for 10 min. Treatment of cells with sodium azide, MβCD and CM (final concentration ×2/3) was performed 30 min before measurements.

To monitor PI(3,4,5)P3 production in response to cAMP stimulation, cells expressing $PH_{CRAC}$-GFP were starved at a density of $5\times10^6$ cells/mL with shaking at 155 rpm for 1 h, followed by 50 nM cAMP pulses every 6 min for 3 h. Cells were collected by centrifugation, washed and resuspended in DB, and introduced into 24-well plates at ~ $2\times10^5$ cells/cm$^2$. To inhibit cell migration, cells were treated with 5 µM Latrunculin A (LatA; Fujifilm Wako, 125-04363) or a mixture of 5 µM LatA and 100 mg/mL MβCD for 10 min. Cells were stimulated with cAMP during time-lapse imaging. All live-cell imaging was performed at 22℃.

**FRAP measurement**

Lipid and membrane protein diffusion was measured by the fluorescence recovery after photobleaching (FRAP) method. An inverted microscope IX83 (Olympus) equipped with a confocal scanning unit (CSU-W1) and a FRAPPA photobleaching module (Andor) was used. Images were acquired every 1 s and photobleaching was performed during image acquisition. The bleached area was 5 µm in diameter for the supported lipid bilayer and 3 µm for the ventral surface of the cell. The fluorescence intensity in the bleached area was divided by the one outside the area to correct for global fading, and normalized by the mean value prior to bleach. The curve was fitted by the following:

$$F(t) = F_o + (F_\infty - F_o)(1 - e^{-t/\tau})$$

where $\tau$, $F_o$, $F_\infty$ are the fitting parameters. Diffusion coefficient (*D*) and mobile fraction ($M_f$) were calculated as $D = \omega^2/4\tau$ and $M_f = (F_\infty - F_o)/(1 - F_o)$ where $\omega$ is radius of the bleached region. The mean value of *D* was calculated if $M_f > 0.2$.

**Cell development on agar plates**

To observe the development of *D. discoideum* (Fig. S7C), $0.6\times10^7$ cells were washed twice and resuspended in DB. Cells were seeded on 1 % DB agar plates containing 0-100 mg/mL MβCD prepared in φ35 mm Petri dishes (IWAKI, 1000-035). After the surface was dried, the plates were placed at 22℃. Images were taken with a stereomicroscope (Olympus, SZX10) equipped with a digital camera (Canon, EOS Kiss X7i).

**Image analysis**

Image analysis was performed using ImageJ, Python and Microsoft Excel. Statistical analysis was performed using Python with a scipy package.

To measure the frequency with which cells formed PI(3,4,5)P3 patches (Fig. 4D), confocal images of cells expressing $PH_{CRAC}$-GFP were taken every 30 s for 15 min. For each cell, the presence or absence of patches was manually scored at each time point during the time course. The frequency of patch formation was calculated by averaging the percentage of frames in which a patch was present over the cell population. Maximum size and duration of patches (Fig. S8A-B) were measured using ImageJ, for which time-lapse images with a shorter 5 s interval were used.

To quantitate PI(3,4,5)P3 production in response to cAMP stimulation, confocal images of cells expressing $PH_{CRAC}$-GFP were taken every 1 s. The average intensity in the cytosol region of each cell was measured and normalized by the average intensity for 1 min before cAMP stimulation. The average and standard deviation for all cells were plotted (Fig. S9). The decrease in intensity from 0 to 30 s after cAMP stimulation was measured for different cAMP concentrations and cell strains, from which dose-response curves were plotted (Fig. 4F-G).

**Table S1. Plasmid list used in this study**

| | Plasmid name | Backbone | Source of reference |
|---|---|---|---|
| 1 | pDM358 act15: mScarlet-I-PldB | pDM358 (hygR) | This study |
| 2 | pDM358 act15: mScarlet-I-PldB$_{N1\text{-}346}$ | pDM358 (hygR) | This study |
| 3 | pDM358 act15: mScarlet-I-PldB$_{N1\text{-}464}$ | pDM358 (hygR) | This study |
| 4 | pDM358 act15: mScarlet-I-PldB$_{PH}$ | pDM358 (hygR) | This study |
| 5 | pDM358 act15: mScarlet-I-PldB$_{PIP2}$ | pDM358 (hygR) | This study |
| 6 | pDM358 act15: mScarlet-I-PldB$_{C887\text{-}1216}$ | pDM358 (hygR) | This study |
| 7 | pDM358 coaA: mScarlet-I-PldB | pDM358 (hygR) | This study |
| 8 | pDM304 coaA: PKBR1$_{N150}$-Neon | pDM304 (neoR) | This study |
| 9 | pDM304 coaA: PKBR1$_{N150}$-K3E-Neon | pDM304 (neoR) | This study |
| 10 | pDM304 coaA: PKBR1$_{N150}$-K7E-Neon | pDM304 (neoR) | This study |
| 11 | pDM304 coaA: PKBR1$_{N150}$-K9E-Neon | pDM304 (neoR) | This study |
| 12 | pDM304 coaA: PKBR1$_{N150}$-K(3,9)E-Neon | pDM304 (neoR) | This study |
| 13 | pDM304 coaA: PKBR1$_{N150}$-K(3,7,9)E-Neon | pDM304 (neoR) | This study |
| 14 | pDM304 act15: cAR1-Neon | pDM304 (neoR) | This study |
| 15 | pDM304 act15: PIKI-Neon | pDM304 (neoR) | This study |
| 16 | pDM1209 act15: PH$_{CRAC}$-GFP | pDM1209 (neoR) | (Honda *et al*., 2021) |

**Table S2. Cell strain list used in this study.**

| | Strain name | Backbone | Source of reference |
|---|---|---|---|
| 1 | AX4 | | Lab stock |
| 2 | AX2 | | Lab stock |
| 3 | *pldB*$^{-}$ | AX2 | (Nagasaki & Uyeda, 2008) |
| 4 | PKBR1$_{N150}$-Neon/AX4 | AX4 | This study |
| 5 | mScarlet-I-PldB/PKBR1$_{N150}$-Neon/AX4 | AX4 | This study |
| 6 | mScarlet-I-PldB$_{N1-346}$/PKBR1$_{N150}$-Neon/AX4 | AX4 | This study |
| 7 | mScarlet-I-PldB$_{N1-464}$/PKBR1$_{N150}$-Neon/AX4 | AX4 | This study |
| 8 | mScarlet-I-PldB$_{PH}$/PKBR1$_{N150}$-Neon/AX4 | AX4 | This study |
| 9 | mScarlet-I-PldB$_{PIP2}$/PKBR1$_{N150}$-Neon/AX4 | AX4 | This study |
| 10 | mScarlet-I-PldB$_{C887-1216}$/PKBR1$_{N150}$-Neon/AX4 | AX4 | This study |
| 11 | cAR1-Neon/AX4 | AX4 | This study |
| 12 | mScarlet-I-PldB/cAR1-Neon/AX4 | AX4 | This study |
| 13 | PIKI-Neon/AX2 | AX2 | This study |
| 14 | mScarlet-I-PldB/PIKI-Neon/AX2 | AX2 | This study |
| 15 | PKBR1$_{N150}$-Neon/AX2 | AX2 | This study |
| 16 | PKBR1$_{N150}$_K3E-Neon/AX2 | AX2 | This study |
| 17 | PKBR1$_{N150}$_K7E-Neon/AX2 | AX2 | This study |
| 18 | PKBR1$_{N150}$_K9E-Neon/AX2 | AX2 | This study |
| 19 | PKBR1$_{N150}$_K(3,9)E-Neon/AX2 | AX2 | This study |
| 20 | PKBR1$_{N150}$_K(3,7,9)E-Neon/AX2 | AX2 | This study |
| 21 | mScarlet-I-PldB/PKBR1$_{N150}$-Neon/AX2 | AX2 | This study |
| 22 | mScarlet-I-PldB/PKBR1$_{N150}$_K3E-Neon/AX2 | AX2 | This study |
| 23 | mScarlet-I-PldB/PKBR1$_{N150}$_K7E-Neon/AX2 | AX2 | This study |
| 24 | mScarlet-I-PldB/PKBR1$_{N150}$_K9E-Neon/AX2 | AX2 | This study |
| 25 | mScarlet-I-PldB/PKBR1$_{N150}$_K(3,9)E-Neon/AX2 | AX2 | This study |
| 26 | mScarlet-I-PldB/PKBR1$_{N150}$_K(3,7,9)E-Neon/AX2 | AX2 | This study |
| 27 | PH$_{CRAC}$-GFP/AX2 | AX2 | This study |
| 28 | mScarlet-I-PldB/PH$_{CRAC}$-GFP/AX2 | AX2 | This study |
| 29 | PKBR1$_{N150}$-Neon/*pldB*$^{-}$ | *pldB*$^{-}$ | This study |
| 30 | mScarlet-I-PldB/PKBR1$_{N150}$-Neon/*pldB*$^{-}$ | *pldB*$^{-}$ | This study |
| 31 | cAR1-Neon/*pldB*$^{-}$ | *pldB*$^{-}$ | This study |
| 32 | PIKI-Neon/*pldB*$^{-}$ | *pldB*$^{-}$ | This study |
| 33 | PH$_{CRAC}$-GFP/*pldB*$^{-}$ | *pldB*$^{-}$ | This study |
| 34 | mScarlet-I-PldB/PH$_{CRAC}$-GFP/*pldB*$^{-}$ | *pldB*$^{-}$ | This study |

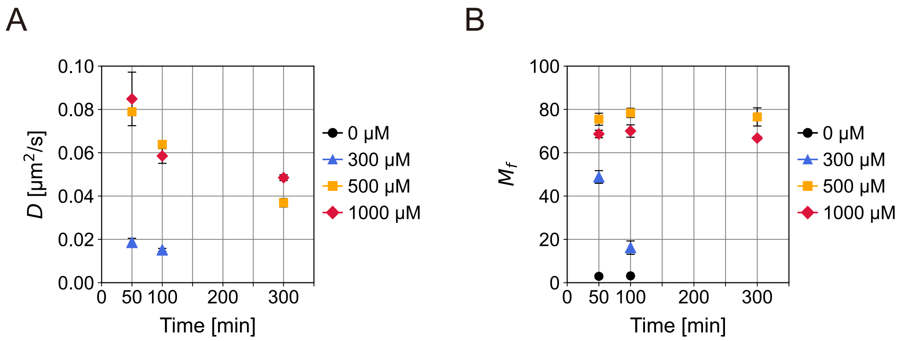


**Figure S1. FIPI treatment of cell extracts maintained membrane fluidity.**
Diffusion coefficients (**A**) and mobile fraction (**B**) of rhodamine-DMPE in DOPC membranes during incubation with FIPI-treated cell extracts (mean ± s.e., N = 2 ~ 5 per condition). Legends, FIPI concentration.

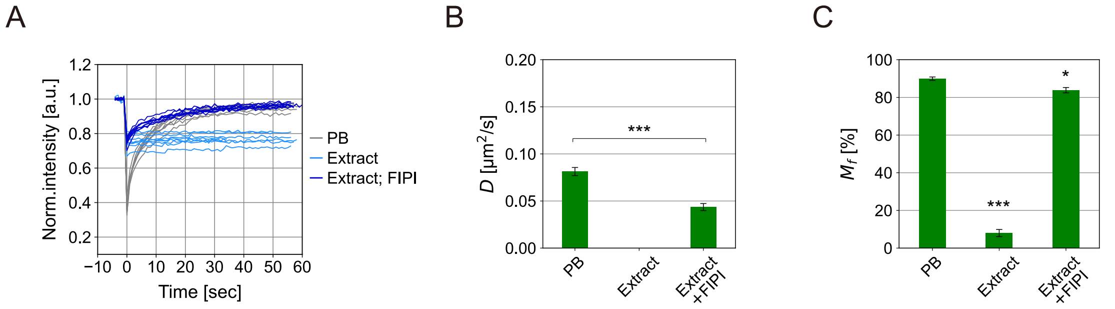


**Figure S2. Loss of membrane fluidity occurred independent of lipid probe.**

(**A)** FRAP curves acquired on membranes containing 18:1-12:0 NBD PC as fluorescent probe. Lipid films were incubated for 40 min with phosphate buffer (PB), cell extracts and extracts treated with 500 µM FIPI (N = 8 each). (**B**) Diffusion coefficients (mean ± s.e.). No value for crude extracts as $M_f$ for all curves was below the 20% criterion. ***$p < 0.001$, Student's t-test. (**C**) Mobile fraction (mean ± s.e.). *$p < 0.05$ and ***$p < 0.001$ compared to PB, one-way ANOVA followed by Kramer-Tukey's post-hoc test.

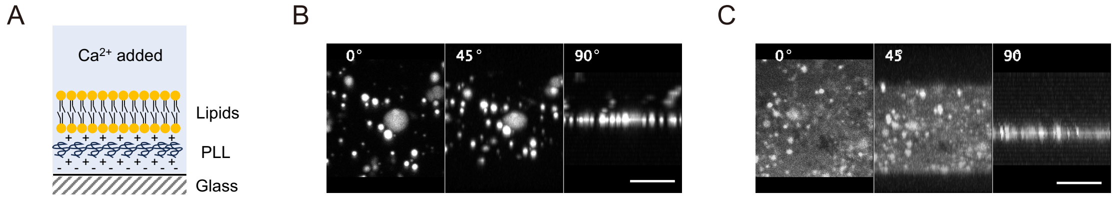


**Figure S3. Lipid films on PLL-coated glass.**

(**A**) Schematic of lipid films on PLL-coated glass. (**B-C**) Lipid films consisting of DOPC and DOPA (mixed at 2:8) formed on PLL-coated coverslips. Maximum intensity projections of Zstacks taken after incubation at pH 6.5 (**B**) and 10.0 (**C**). Numbers in images, projection angle. Scale bars, 5 μm.

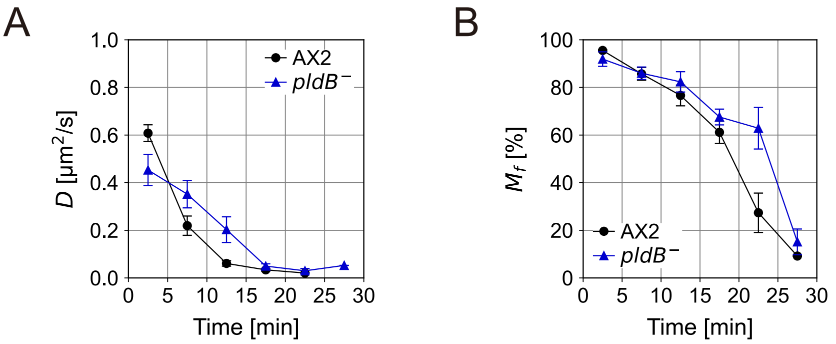


**Figure S4. The *pldB⁻* extract had the same effect on membrane fluidity as that of wild-type cells.**

Diffusion coefficients (**A**) and mobile fraction (**B**) of rhodamine-DMPE in DOPC membranes during incubation with AX2 and *pldB⁻* extracts (mean ± s.e., N = 2 ~ 9 per plot).

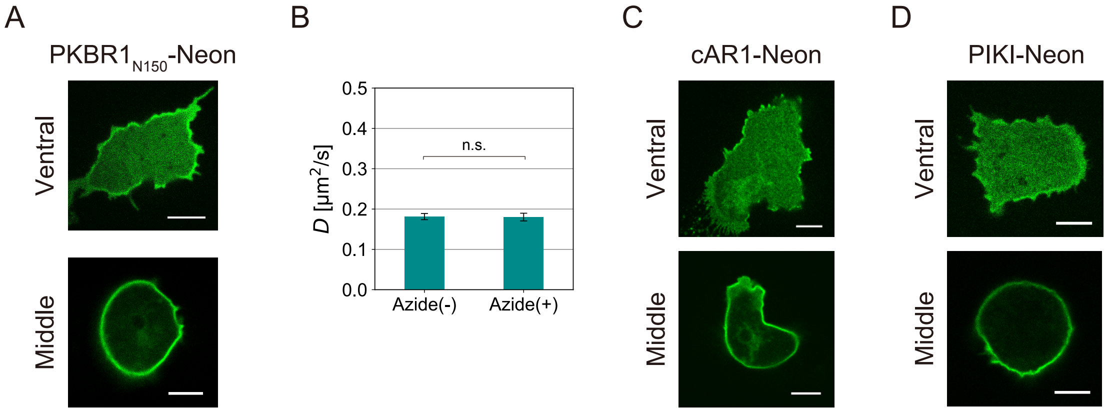


**Figure S5. Confocal images of membrane proteins used for FRAP measurements.**
(**A**) PKBR1$_{N150}$-Neon. (**C**) cAR1-Neon. (**D**) PIKI-Neon. Images were taken at the ventral surface (*top*) and the middle plane (*bottom*). Scale bars, 5 μm. (**B**) Diffusion coefficients (mean ± s.e.) of PKBR1$_{N150}$-Neon in AX4 treated with 1 mM sodium azide (N = 15 cells each). n.s., Not significant, Student's t-test.

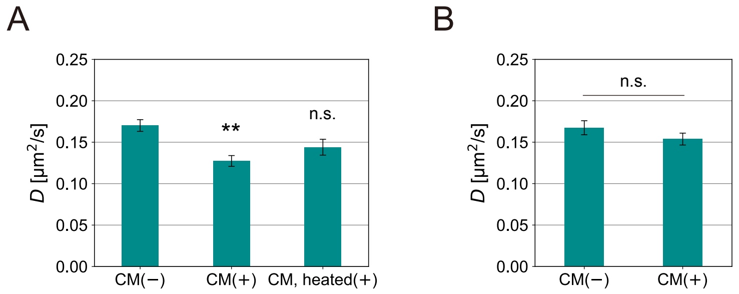


**Figure S6**. **Cellular secretions affect the diffusion rate of PKBR1 through PldB.**

Diffusion coefficients (mean ± s.e.) measured by FRAP. (**A**) PKBR1$_{N150}$-Neon in AX4 cells treated with DB (mock control), conditioned medium (CM) and heat-treated CM (N = 20 cells each). ***p* < 0.01 and n.s.; not significant, compared to mock control, one-way ANOVA followed by Kramer-Tukey's post-hoc test. (**B**) PKBR1$_{N150}$-Neon in *pldB*$^{-}$ treated with DB and CM (N = 30 and 27 cells). Student's t-test.

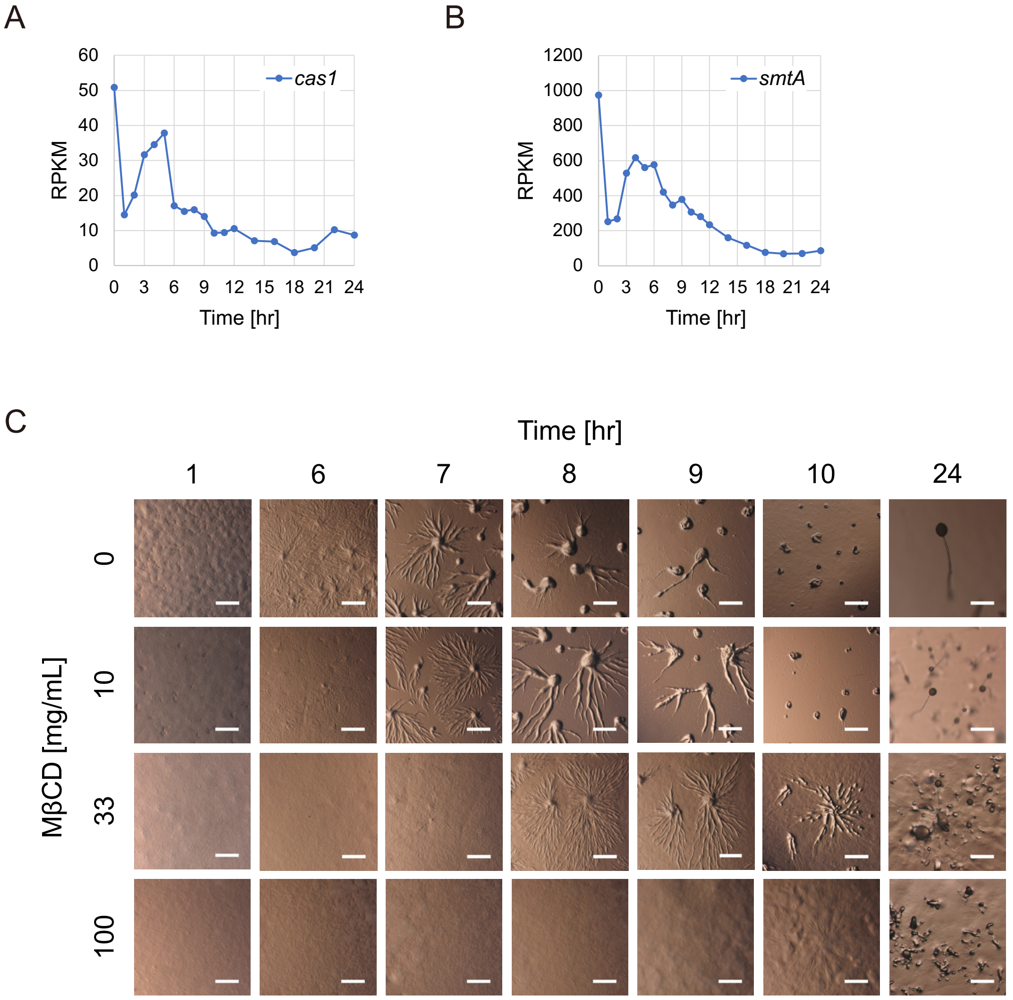


**Figure S7**. **Function of sterols in *Dictyostelium discoideum* development.**
(**A-B**) Time course of expression of genes responsible for sterol synthesis. (**A**) *cas1*. (**B**) *smtA*. Plotted data are from publicly available RNA-seq (3, 4). (**C**) Snapshots of AX2 cells developed on agar plates containing MβCD at the concentrations indicated on the left. Scale bars, 1 mm (1-10 hr) and 300 μm (24 hr).

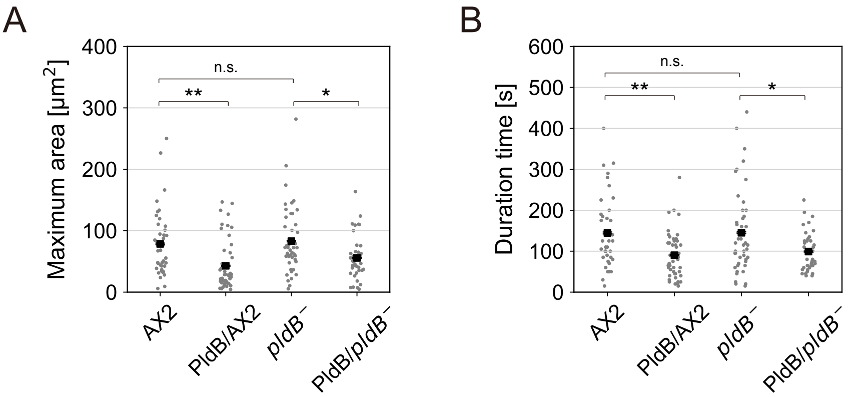


**Figure S8. PldB reduces the size and duration of PIP3 patches.**

(**A**) Maximum expansion area and (**B**) duration of $PH_{CRAC}$-GFP patches in AX2, AX2 over-expressing mScarlet-I-$PldB_{FL}$ (PldB/AX2), *pldB*$^{-}$ and *pldB*$^{-}$ over-expressing mScarlet-I-$PldB_{FL}$ (PldB/*pldB*$^{-}$) (mean ± s.e., N = 39, 47, 46 and 38 patches). Each dot corresponds to an individual patch. *$p < 0.05$, **$p < 0.01$. and n.s.; not significant, one-way ANOVA followed by Kramer-Tukey's post-hoc test.

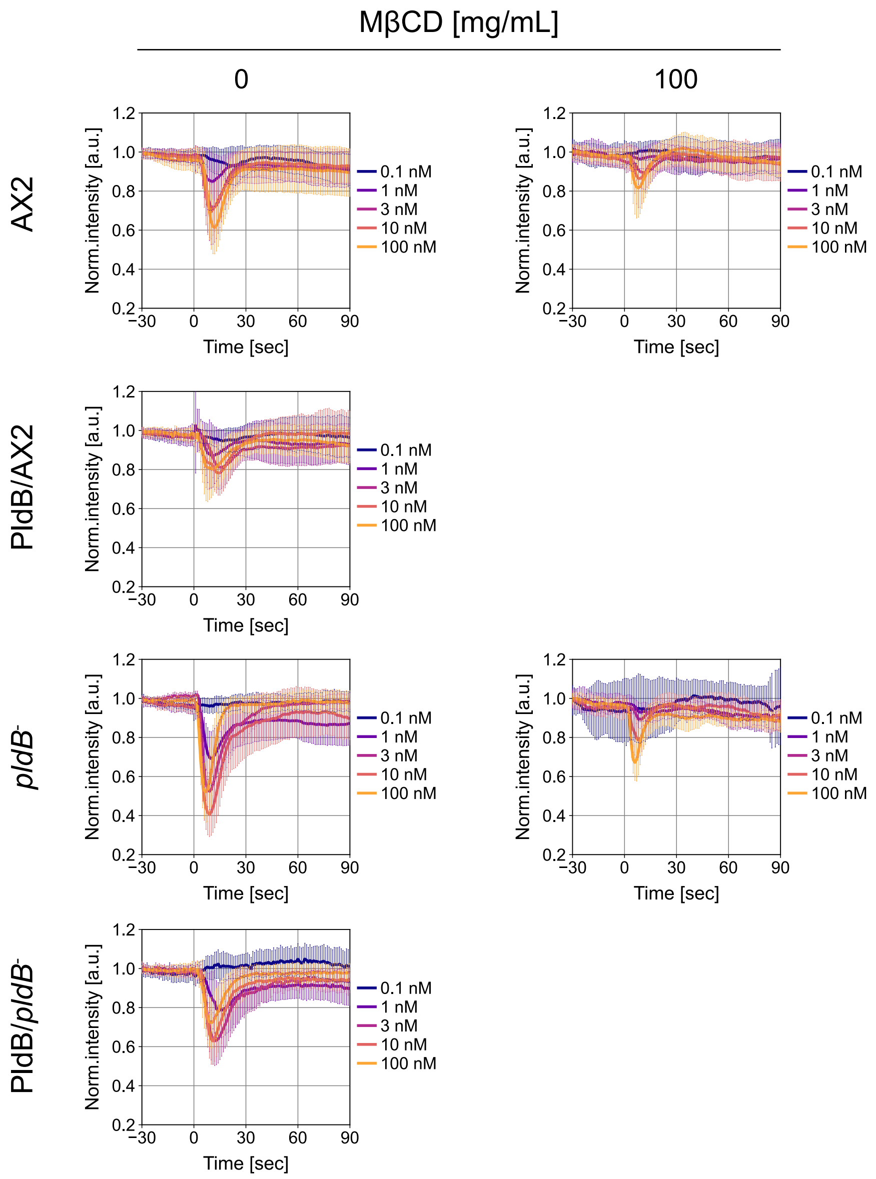


**Figure S9. PldB suppresses PI(3,4,5)P3 production in response to cAMP stimulation.** Time profiles of $PH_{CRAC}$-GFP intensity at cytosol (mean ± s.d.). cAMP was uniformly added at $t$ = 0 s. Legend, cAMP concentration. Intensity was normalized by the average value before cAMP addition. AX2, AX2 over-expressing mScarlet-I-$PldB_{FL}$ (PldB/AX2), *pldB*⁻ and *pldB*⁻ over-expressing mScarlet-I-$PldB_{FL}$ (PldB/*pldB*⁻) cells were used. Cells were pretreated with 5 µM LatA and MβCD.

**Movie legend**

**Movie S1. DOPC membranes after application of the cell extract.**

Time-lapse confocal images of rhodamine-DMPE mixed into the DOPC membrane. Images were acquired every 15 sec immediately after applying the cytosolic fraction collected from *D. discoideum*. Numbers, time (min). Scale bar, 10 μm.
